# Noise-driven multistability versus deterministic chaos in phenomenological semi-empirical models of whole-brain activity

**DOI:** 10.1101/2020.07.31.231712

**Authors:** Juan Piccinini, Ignacio Perez Ipina, Helmut Laufs, Morten Kringelbach, Gustavo Deco, Yonatan Sanz Perl, Enzo Tagliazucchi

## Abstract

An outstanding open problem in neuroscience is to understand how neural systems are capable of producing and sustaining complex spatiotemporal dynamics. Computational models that combine local dynamics with *in vivo* measurements of anatomical and functional connectivity can be used to test potential mechanisms underlying this complexity. We compared two conceptually different mechanisms: noise-driven switching between equilibrium solutions (modeled by coupled Stuart-Landau oscillators) and deterministic chaos (modeled by coupled Rossler oscillators). We found that both models struggled to simultaneously reproduce multiple observables computed from the empirical data. This issue was especially manifest in the case of noise-driven dynamics close to a bifurcation, which imposed overly strong constraints on the optimal model parameters. In contrast, the chaotic model could produce complex behavior over an ampler range of parameters, thus being capable of capturing multiple observables at the same time with good performance. Our observations support the view of the brain as a non-equilibrium system able to produce endogenous variability. We presented a simple model capable of jointly reproducing functional connectivity computed at different temporal scales. Besides adding to our conceptual understanding of brain complexity, our results inform and constraint the future development of biophysically realistic large-scale models.

The quote *“What I cannot create, I do not understand”* was found written in the blackboard of celebrated physicist Richard Feynman at the time of his death. This sentence suggests a way forward for neuroscientists interested in unravelling the principles behind the richness and complexity of spontaneous brain dynamics. Over the last decades, tremendous advances in neuroimaging enabled the construction of whole-brain activity models with real predictive power in the statistical sense. It is now possible to *create* realistic complex dynamics, instead of passively screening for their presence in neuroimaging data. We contrasted two different types of building blocks (i.e. two choices of local dynamics) and tested their capacity to reproduce the empirical data, with the purpose of increasing our conceptual understanding of the mechanisms behind large-scale spontaneous activity in the human brain.

## I. INTRODUCTION

Since the mid 1990s, the spontaneous large-scale^1^ dynamics of the human brain have attracted a growing body of research^2^. Previously disregarded as experimental noise, it became increasingly clear that the fluctuating dynamics of endogenous brain activity were not random, but highly structured in the spatial and temporal domains^3,4^. During rest, this activity self-organizes into recurrent patterns that overlap with different functional systems of the brain, known as *resting state networks* (RSN)^5^. Since the activity patterns evoked by cognitive tasks and sensory stimulation can be put into correspondence with the RSN^6^, it has been proposed that the spontaneous dynamics of the brain reflect a metastable exploration of states that facilitate rapid reactivity upon environmental demands^3,7–9^. According to this view, evolution has shaped the brain as an itinerant dynamical system which is always close to configurations associated with sensory, cognitive or motor functions^10^.

The complex spatiotemporal organization of large-scale neural activity is a robust finding demonstrated across several mammalian species^11–16^ using imaging tools covering an ample range of spatial and temporal scales, including functional magnetic resonance imaging (fMRI)^5^, electroencephalography (EEG)^17^, magnetoencephalography (MEG)^18^, electrocorticography (ECoG)^19^, functional near-infrared spectroscopy (fNIRS)^20^, as well as some of their combinations^21–23^. The spatial and temporal properties of resting brain activity depend on the global state of consciousness^24–27^, and present alterations specific to several neurological and psychiatric conditions^28^, suggesting their potential application as disease biomarkers in precision medicine^29^. In spite of their obvious relevance for basic and clinical neuroscience, the mechanisms supporting the emergence of complexly patterned spontaneous brain activity remain, to a large extent, unknown.

Computational models show promise to identify some of these mechanisms^8^. Modern neuroimaging techniques are capable of mapping large-scale anatomical and functional connectivity with unprecedented breadth and accuracy^30^, which can be leveraged to construct whole-brain activity models that reproduce statistical features of empirical data^31^. In these models, local dynamics are coupled by the anatomical connectivity of the brain, and model parameters are then optimized to reproduce certain observables of interest computed from neuroimaging recordings^32^. By exploring the consequences of changing the local dynamics and global coupling, these semi-empirical^33^ models can inform the mechanisms underlying the features of spontaneous brain activity represented in the chosen observable. Models of whole-brain activity thus constructed have been intensively explored in the past few years to investigate the mechanisms underlying healthy and pathological brain states, to simulate the consequences of externally induced brain stimulation, and with the purpose of data augmentation for machine learning classification of neuroimaging data^34–45^.

The choice of local dynamics determines the range of qualitatively distinct behaviors of the model, its complexity (i.e. number of free parameters), and in turn depends on the desired level of biophysical realism^32,46^. When it comes to modeling the features of spontaneous brain activity recorded with techniques such as fMRI, EEG and MEG, whole-brain models emerge as the natural choice. A common property of phenomenological^47^ models is the inclusion of noise-driven dynamics near a bifurcation to reproduce certain key features of spontaneous brain activity (e.g. metastability, organization into RSN)^7,39,48,49^. For deterministic models attracted to stable or periodic solutions, noise is fundamental to avoid that dynamics become stuck in a state of equilibrium. Thus, the *ad hoc* introduction of noise in a dynamical system near a bifurcation ensures stochastic switching between different attractors, endowing the simulation with the kind of variability seen in the empirical data.

Another key choice to be made when constructing a whole-brain model is the observable to be reproduced. Since the temporal evolution of complex systems such as the human brain is considered difficult or outright impossible to predict, a meaningful observable should encode the behavior of the system in a statistical sense. Two well-known examples are the functional connectivity (FC) matrix, containing all pairwise correlations between the local activity time series^43,50^, and the functional connectivity dynamics (FCD), which represents the similarity between FC matrices computed over short windows at different times^39,49^. The first of these two observables is useful to represent functional coupling over long or “static” time scales, while the second aims to capture the dynamic switching between metastable states^9^. Other observables can be defined, either assessing static or dynamic characteristics of brain activity.

The possibility of defining multiple meaningful observables raises a problem in the process of constructing a whole-brain activity models based on noise-driven equilibrium dynamics. If the system must be poised at a particular combination of parameters (i.e. bifurcation) to ensure the reproduction of a certain observable, does it follow that the same combination of parameters will successfully reproduce other equally meaningful observables? An affirmative answer seems unlikely.

Since a systematic evaluation of how noise-driven equilibrium models perform when simultaneously reproducing multiple observables is lacking, severe limitations could exist for this class of models to explore mechanisms involving more than one feature of spontaneous brain activity. These potential limitations prompt the need to investigate models that are capable of producing complex brain dynamics over a wide range of parameter values.

In this work we explore deterministic chaos as an alternative to noise-induced multistability to reproduce statistical observables computed from resting state fMRI recordings. Chaotic dynamics unfold in the proximity of strange attractors, i.e. complicated fractal sets that give rise to bounded but non-periodic trajectories that are highly dependent on the initial conditions^51^. Among others, mechanisms such as heteroclinic cycling^52^ and chaotic itineracy^53^ endow networked chaotic dynamical systems with complex metastable dynamics in the absence of noise. While the possibility of intrinsically chaotic brain dynamics has been explored in neural recordings acquired from several model organisms investigated at an ample range of spatial and temporal scales^54–58^, the consequences of including deterministic chaos in the local dynamics of semi-empirical whole-brain activity models remains comparatively unexplored. Our purpose is not to engage in the long-standing discussion of deterministic chaos as a fundamental property of brain activity^59,60^. Instead, we are interested in the more pragmatic question of whether deterministic chaos results in models capable of avoiding some limitations intrinsic to noise-driven equilibrium dynamics. In particular, we are interested in how both types of models perform when attempting to simultaneously reproduce multiple empirical observables. Our main motivation is to identify the simplest dynamics capable of reproducing several key features of spontaneous brain activity at the same time, with the purpose of informing the future development of more biophysically realistic models^61^.

For this purpose, we investigated the dynamics of Stuart-Landau^39^ and Rossler^62^ oscillators coupled by an anatomical connectivity network obtained *in vivo* from diffusion tensor imaging (DTI)^63^. These dynamical systems constitute simple examples capable of exhibiting noise-driven multistability and deterministic chaos, respectively. The Stuart-Landau oscillator undergoes a supercritical Hopf bifurcation, thus capturing the dichotomy between fixed-point noisy dynamics and harmonic oscillatory activity^39^. Due to its conceptual simplicity, it has been used as the basis of phenomenological models in several recent publications^37,38,40,42–44^. For certain parameters, the Rossler system gives rise to a strange attractor with a single manifold, and is considered one of the simplest examples of chaotic dynamics^62^. We explored and compared how well these two models reproduced several common observables computed from fMRI recordings.

## II METHODS

### A. Participants and EEG-fMRI data acquisition

A cohort of 63 healthy subjects participated in the original experiment (36 females, mean ± SD age of 23.4 ± 3.3 years). Written informed consent was obtained from all subjects. The experimental protocol was approved by the local ethics committee (Goethe-Universität Frankfurt, Germany, protocol number: 305/07). The subjects were reimbursed for their participation. All experiments were conducted in accordance with the relevant guidelines and regulations, and the Declaration of Helsinki.

Participants were scanned for 50 minutes using previously published acquisition parameters^26^. From the group of 63 subjects we selected a subgroup of 9 participants who remained awake throughout the complete duration of the scan (confirmed by assessment of the simultaneous EEG^64^). In this way, we obtained unusually long fMRI recordings with the purpose of robustly estimating observables related to FC dynamics.

### B. fMRI data preprocessing

Using Statistical Parametric Mapping (SPM8, www.fil.ion.ucl.ac.uk/spm), raw fMRI data were realigned, normalized and spatially smoothed using a Gaussian kernel with 8 mm^3^ full width at half maximum. Data was then resampled to 4 × 4 × 4 mm resolution. Note that re-sampling introduced local averaging of blood-oxygen-level-dependent (BOLD) signals, which were eventually averaged over larger cortical and sub-cortical regions of interest as determined by the automatic anatomic labeling (AAL) atlas^65^. Data was denoised by regressing out cardiac, respiratory and residual motion time series^66^, and then band-pass filtered in the 0.01–0.1 Hz range using a sixth order Butterworth filter^67^.

### C. Anatomical connectivity matrix

The anatomical connectivity matrix was obtained applying diffusion tensor imaging (DTI) to diffusion weighted imaging (DWI) recordings from 16 healthy right-handed participants (11 men and 5 women, mean age: 24.75 ± 2.54 years) recruited online at Aarhus University, Denmark. Subjects with psychiatric or neurological disorders (or a history thereof) were excluded from participation. We refer to a previous publication for details of the MRI acquisition parameters^43^.

Anatomical connectivity networks were constructed following a three-step process. First, the regions of the whole-brain network were defined using the AAL atlas. Second, the connections between nodes in the whole-brain network (i.e. edges) were estimated applying probabilistic tractography to the DTI data obtained for each participant. Third, results were averaged across participants.

DTI preprocessing was performed using the *probtrackx* tool of the FSL diffusion imaging toolbox (Fdt) (www.fsl.fmrib.ox.ac.uk/fsl/fslwiki/FDT) with default parameters. Next, the local probability distributions of fiber directions were estimated at each voxel. The connectivity probability from a seed voxel *i* to another voxel *j* was defined as the proportion of fibers passing through voxel *i* that reached voxel *j*, sampling a total of 5000 streamlines per voxel. This was extended from the voxel to the region level, i.e. in a region of interest consisting of n voxels, 5000 × n fibers were sampled. The connectivity probability from region *i* to region *j* was calculated as the number of sampled fibers in region *i* that connected the two regions, divided by 5000 × n, where n represents the number of voxels in region *i*. The resulting anatomical connectivity matrices were thresholded at 0.1% (i.e. a minimum of five streamlines), resulting in the anatomical connectivity matrices *C_i,j_* used in the models.

### D. Whole-brain model construction

The general procedure to construct whole-brain models is presented in Fig.1. Following previous work, we constructed computational models of whole-brain activity by assigning local dynamical rules to 90 nodes spanning the whole cortical and subcortical grey matter. These nodes were coupled using an anatomical connectivity matrix *C_i j_* which contained in its *i, j* entry an estimate of the number of white matter tracts connecting nodes *i* and *j* (see previous section). We introduced a parameter *G* to globally scale the *C_i j_*, thus modeling changes in the overall strength of inter-areal coupling.

To determine the natural frequency of the local dynamics, we estimated the most dominant frequency at each node (*ω_j_*) as the peak of the Fourier-transformed fMRI time series (averaged across all participants). Consistent with previous reports^39,67^, these frequencies were in the 0.04-0.07 Hz range.

The fMRI signal corresponding to node *j* was simulated by the variable *x_j_* of the differential equation modeling the local dynamics, integrated using a second-order Runge–Kutta algorithm with a time step of 0.1. For each model and each parameter combination we produced a number of simulations matching the number of subjects in the dataset. Simulated time series were downsampled to match the sampling frequency of the fMRI data.

#### 1. Hopf model

For noise-driven multistability we considered local dynamics given by the Stuart-Landau oscillator, which is equivalent to the normal form of a supercritical Hopf bifurcation. The differential equations for node *j* are given by,

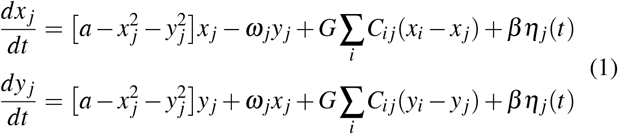

where *η_j_*(*t*) represents additive Gaussian noise, *β* is the noise scaling parameter, *G* is the anatomical connectivity scaling parameter, and *a* is the bifurcation parameter. For an uncoupled Hopf bifurcation, *a* < 0 results in a fixed-point attractor and *a* > 0 in an attracting limit cycle, leading to harmonic oscillations at the natural frequency of the node *ω_j_* (estimated from the empirical data). For *a* ≈ 0 the system switches between both solutions as a consequence of the additive noise term, as shown in Fig. 1 and Eq. 1.

**FIG. 1.**
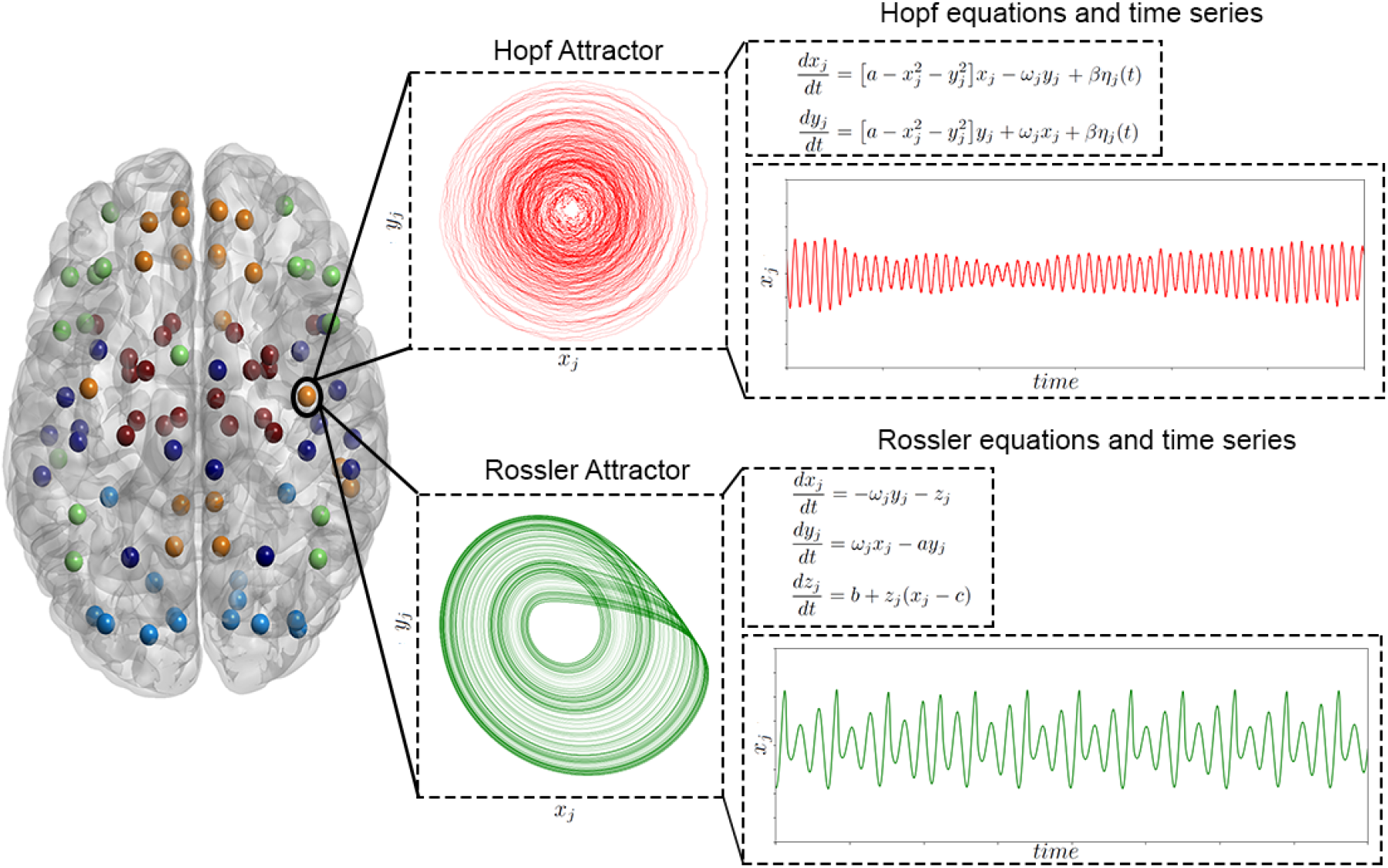
Schematic of the process followed to construct the whole-brain models. The 90 nodes spanning the whole cortical and subcortical grey matter were coupled by the anatomical connectivity matrix *C*_*i, j*_ estimated using probabilistic tractography applied to DTI data. We constructed two models differing in the choice of local dynamics: a model presenting noise-driven multistability (Stuart-Landau oscillators, red), and a model presenting local deterministic chaos (Rossler oscillators, green). In both cases, local dynamics were coupled according to the *C*_*i, j*_ matrix scaled by a factor *G*. On the right, we illustrate typical 2D phase space projections and time series for the Stuart-Landau oscillator near the Hopf bifurcation, and for the Rossler oscillator at the chaotic regime. In the first case, dynamics present complex amplitude modulations due to noise-induced switching between the fixed-point and the limit cycle attractors. In the second case dynamics present aperiodic behavior unfolding in the proximity of a strange attractor with a single manifold.

#### 2. Rossler model

For deterministic chaos we considered local dynamics given by the Rossler oscillator, which is a system of three non-linear ordinary differential equations exhibiting chaotic dynamics for certain parameter combinations. The differential equations for node *j* are given by,

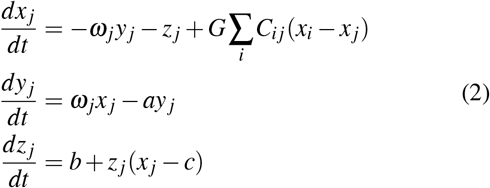

where *a*, *b* and *c* are free model parameters and *G* is the global scaling factor of the anatomical connectivity. We fixed parameters *b* = 0.01 and *c* = 13.44 using a genetic algorithm (see the following section). Since the natural frequencies of brain activity are *ω*_0_ ~ 0.3, the equations were rescaled using a factor *γ* = 0.3 to match the empirical frequencies, while preserving the dynamics of the oscillators. We note that the chosen parameters lead to chaotic dynamics (see Figure 1).

### E. Rossler parameter selection

Since there is a mismatch in the number of free parameters between models, we applied a stochastic optimization procedure (genetic algorithm) to fix two of the three parameters of the Rossler model. Following previous work^43^, we used 1 minus the structural similarity index (1-SSIM) as the distance metric between simulated and empirical FC matrices^68^. The SSIM is defined as 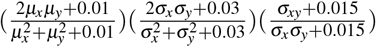, where *x* and *y* stand for the simulated and average empirical FC, and *µ_x_*, *µ_y_*, *σ_x_*, *σ_y_* and *σ_xy_* correspond to the local means, standard deviations, and co-variances of FC matrices *x* and *y*, respectively^69^. The SSIM simultaneously factors the correlation and Frobenius distances between matrices, and can be intuitively understood as an intermediate between both^68^.

The genetic algorithm started with a generation of 10 sets of parameters (“individuals”) chosen randomly in the range [0.0, 0.3], [0.0, 0.3], [0.0, 14] and [0.0, 3.0] for *a*, *b*, *c* and *G*, respectively. A score proportional to the target function was assigned to each individual. Afterwards, a group of individuals was chosen based on their score (“parents”), and operations of crossover, mutation and elite selection were applied to them to create the next generation. These three operations can be briefly described as follows: 1) elite selection occurs when an individual of a generation shows an extraordinarily low target function in comparison to the other individuals, thus this solution is replicated without changes in the next generation; 2) the crossover operator consists of combining two selected parents to obtain a new individual that carries information from each parent to the next generation; 3) the mutation operator can change one selected individual to induce a random alteration.

Following our previous work, 20 % of each new generation was created by elite selection and 80 % by crossover of the parents, with a 5 % chance of possible mutations within this last group. A new population was thus generated (“offspring”) and used iteratively as the next generation until the halting criteria were met, consisting of either reaching an average target function less than 10^−5^ across the last 50 generations, or obtaining a constant value for the target function during 50 generations. After applying the optimization algorithm, the parameter values corresponding to the best fit were *a* = 0.3, *b* = 0.01, *c* = 13.44 and *G* = 1.5. This procedure was used to fix parameters *b* and *c*, while parameters *a* and *G* were explored in the same way as for the Hopf model.

### F. Phase synchrony and metastability

We extracted the phases of the band-pass filtered fMRI signals for each of the 90 brain regions and for each subject. The phases were obtained by applying the Hilbert transform to the filtered time series, which resulted in the associated analytic narrowband signal, *a*(*t*). The analytic signal *a*(*t*) of a signal *x*(*t*) is defined as *a*(*t*) = *x*(*t*) + *iH*[*x*(*t*)], where *i* is the imaginary unit and *H*[*x*(*t*)] denotes the Hilbert transform of *x*(*t*).

We quantified the global degree of synchrony between the nodes across time with the Kuramoto order parameter, *R*(*t*), a measure of phase locking given by:

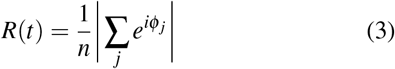

where *n* is the total number of nodes and *ϕ_j_*(*t*) the instantaneous phase of the narrowband signal at node *j*^44^. Thus, the Kuramoto order parameter measures the modulus of the average phase of the system at each time point and takes values from 0 to 1. Here, 0 represents complete absence of phase synchrony and 1 indicates full synchronization.

We calculated the temporal average and standard deviation of the Kuramoto order parameter per subject and subsequently averaged these measures across subjects. We termed these observables *synchrony* and *metastability*, respectively. The synchrony represents the global, temporally averaged degree of synchronization between all the nodes in the system, whereas the metastability gives information about temporal variability in the level of synchronization.

### G. Model fitting

For each distance metric to be optimized, we performed an exhaustive parameter space exploration by varying the two free parameters of the models, corresponding to the global coupling parameter *G* and the free model parameter *a*, where *a* was changed homogeneously across all nodes. The global coupling strength *G* was varied from 0 to 3 in steps of 0.05, the Hopf bifurcation parameter *a* from 0.15 to 0.15 (range chosen to include the bifurcation point), and the Rossler parameter *a* from 0.01 to 3.1 (range chosen to ensure positive Lyapunov exponents of the local dynamics), both in steps of 0.005. For each parameter combination we computed multiple observables to be defined in the next section; subsequently, we used different distance metrics to compare the empirical and simulated observables. This procedure was performed nine times under exactly the same conditions, and the resulting distance metrics were then averaged, selecting the optimal parameters as those that yielded the lowest value of the averaged distance metric.

#### 1. Functional connectivity matrix

We estimated the functional connectivity (FC) matrix for each parameter combination by computing the Pearson correlation coefficient between fMRI signals from all pairs of regions of interest. Subsequently, we calculated the distance between the empirical and simulated FC matrices using three different metrics: Frobenius distance (normalized by the norm of the empirical FC matrix), correlation distance (1-Correlation) and 1-SSIM (defined in a previous section). This resulted in one distance metric value for each parameter combination and per each simulation.

#### 2. Comparison between empirical and simulated synchrony and metastability

The phase synchrony and metastability were computed for each choice of parameters from the filtered simulated time series by applying the same procedure as for the empirical data.

We computed the difference between the simulated and empirical observables, and then normalized the results dividing by the empirical value:

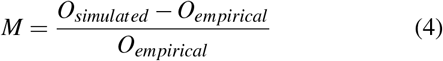

where *M* represents the distance metric, *O_simulated_* denotes the synchrony or metastability obtained from the simulated data, and *O_empirical_* denotes the empirical values.

#### 3. Functional connectivity dynamics

In order to characterize the time-dependent structure of the resting state fluctuations, we computed the FCD matrices^46,49^. Each full-length BOLD signal of 50 minutes was split up into *M* = 148 sliding windows of 60 seconds, overlapping by 40 seconds. For each sliding window centered at time *t*, we calculated an FC matrix, FC(*t*). The FCD is a *M* × *M* symmetric matrix whose (*t*_1_, *t*_2_) entry is defined by the Pearson correlation coefficient between the upper triangular parts of the two matrices FC(*t*_1_) and FC(*t*_2_). FCD matrices were computed for each of the nine participants and simulations, also exhaustively exploring the model parameters. In order to capture fluctuations in correlations across the whole signal spectrum, the time series were not filtered when evaluating this observable.

To compare FCD matrices we collected their upper triangular elements (across all participants) and compared the resulting empirical distribution with that obtained across all simulations using the Kolmogorov-Smirnov (KS) statistic as a distance metric. This metric quantifies the maximum difference between the cumulative distribution functions of the two samples.

## III RESULTS

### A. Group-averaged empirical observables

We computed a set of statistical observables from empirical resting state fMRI data acquired during long scanning sessions (50 minutes) for 9 healthy awake participants. For this purpose, the blood-oxygen-level-dependent (BOLD) signals corresponding to each voxel were averaged over 90 cortical and sub-cortical regions of interest, determined by the automatic anatomic labeling (AAL) atlas^65^. We obtained the group-level functional connectivity (FC) matrix by computing all pairwise correlations between the BOLD signals corresponding to the 90 regions of interest, averaging afterwards the FC matrices across participants (Fig. 2).

**FIG. 2.**
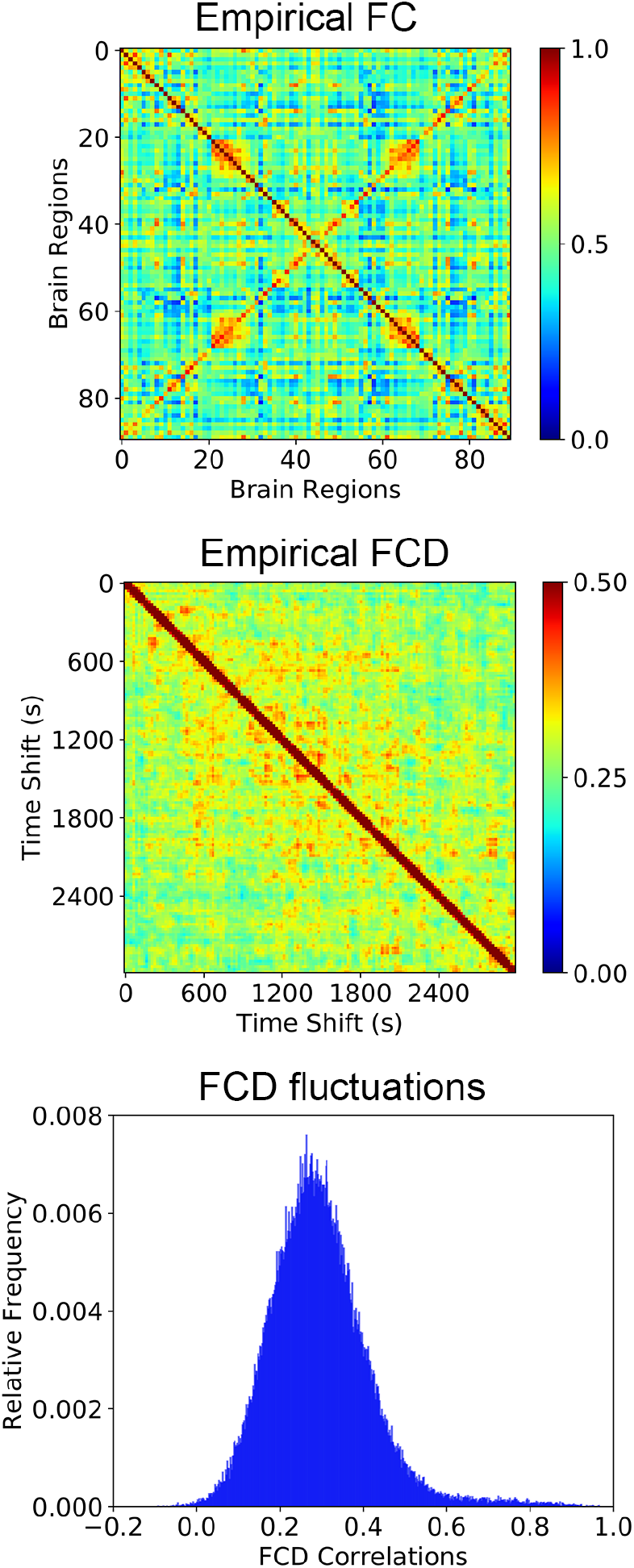
Average empirical static and dynamic observables. The average FC matrix (top), an example FCD matrix (center), and the histogram of FCD values (bottom) computed from the empirical fMRI data.

FCD matrices were obtained by computing the correlation between FC matrices corresponding to sliding windows of 60 seconds with a 40 seconds overlap. Fig.2 displays the FCD matrix and the histogram obtained from the upper-diagonal part of the matrix (pooled across all participants). Other empirical observables related to the dynamics of brain activity, such as phase synchrony and metastability^44^ were computed for each participant and then averaged across the whole sample, following the procedure outlined in the Methods section.

### B. Fitting whole-brain models to empirical FC matrices

We performed an exhaustive exploration of the parameter space for the Hopf and Rossler models, computing each of the three different metrics outlined in the Methods section (1-Correlation, 1-SSIM, and Frobenius distance) for each combination of parameter values. The results (averaged across all realizations) are shown in Fig.3A.

**FIG. 3.**
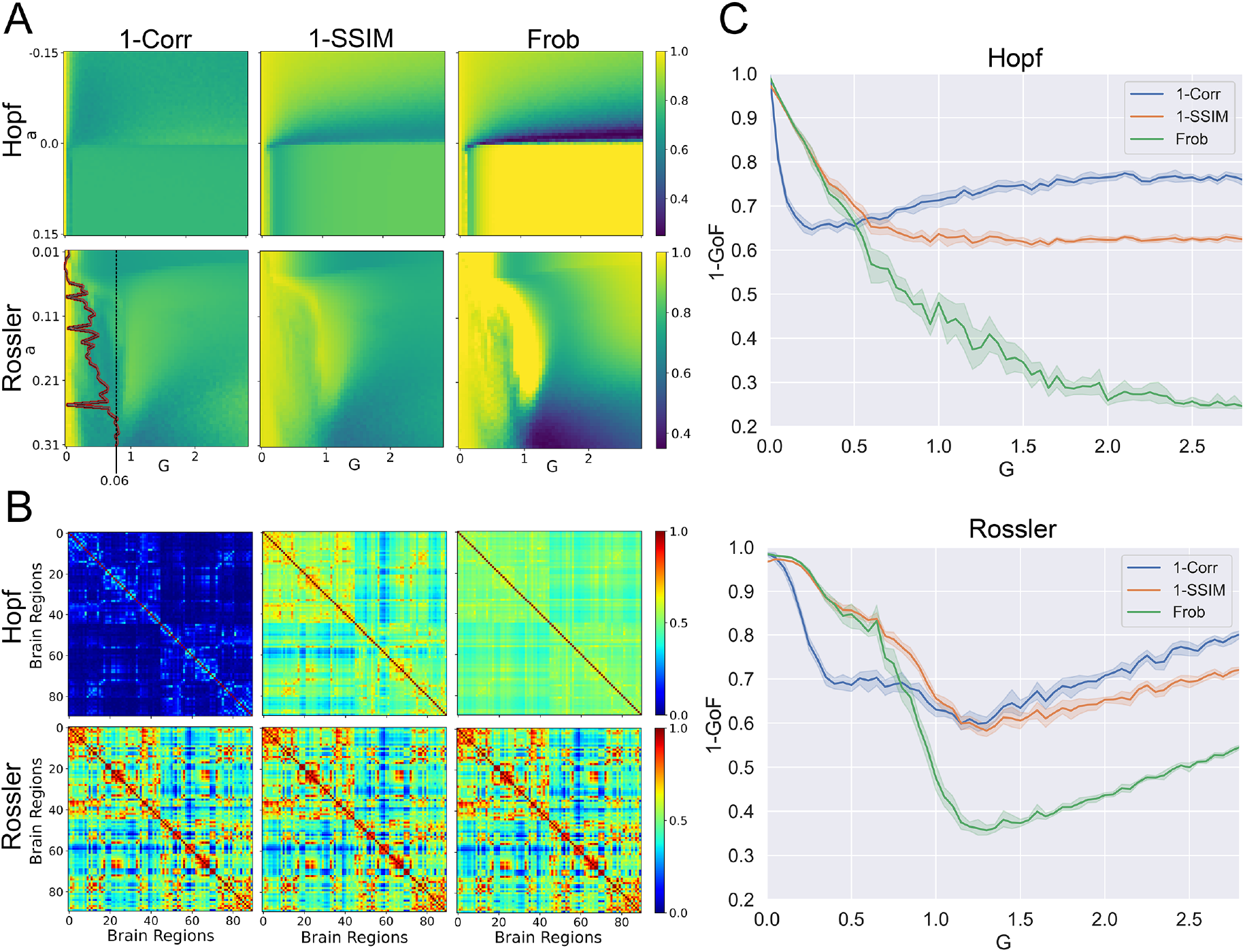
Whole-brain models fitted to static FC metrics. (a) Exhaustive exploration of the 2D parameter space for the Hopf and Rossler models. The color-coded matrices display the distance between simulated and empirical FC matrices according to three different metrics: 1-Correlation, 1-SSIM, and Frobenius distance. The Lyapunov exponent for the local dynamics of the Rossler model is shown as a red curve superimposed over the 1-Correlation matrix as a function of parameter *a*. (b) The FC matrices computed using the optimal parameter combinations obtained from each distance metric. (c) The three different metrics computed for both models as a function of the coupling parameter *G*, using the best parameter *a* obtained from optimizing the Frobenius distance.

As expected from previous work^44^ we observed that for the Hopf model the best fit occurred close to the bifurcation point for all metrics. For the Rossler model, instead, comparatively good fits were obtained for the three metrics across an ample range of parameter values. The local dynamics of the Rossler model presented positive Lyapunov exponents across the explored range of parameters (shown as the red curve superimposed over the 1-Correlation panel).

Figure 3B shows the FC matrices obtained for the optimal parameter combinations using each metric as target function. We observed higher consistency in the FC matrices simulated with the Rossler model. Further characterization of these results is shown in Fig. 3C, presenting 1-GoF of the metrics for the model parameter *a* optimized according to the Frobenius distance, as a function of the scaling parameter *G*. An optimal coupling parameter could be found for all metrics, with the exception of the Frobenius distance and 1-Correlation in the Hopf model. For the Rossler model, all metrics presented a clearly defined optimal *G* value, which was also comparatively similar between them.

### C. Fitting whole-brain models to dynamic observables

Next, we performed the same analysis but with the purpose of reproducing three dynamic observables: the distribution of FCD values, the synchrony, and the metastability. The left panel of Fig. 4A shows the KS distance between the distribution of simulated and empirical FCD values for all parameter combinations. Once more, we can see that the Rossler model presented an ample region of optimal values. The right panel of the figure shows the histograms of FCD values obtained for the optimal parameters optimized using the KS distance, along with the empirical distribution in blue. We can see that the optimal FCD distribution for the Rossler model appears to be skewed towards the right, while optimal distribution for the Hopf model follows a normal distribution centered around the empirical mean.

**FIG. 4.**
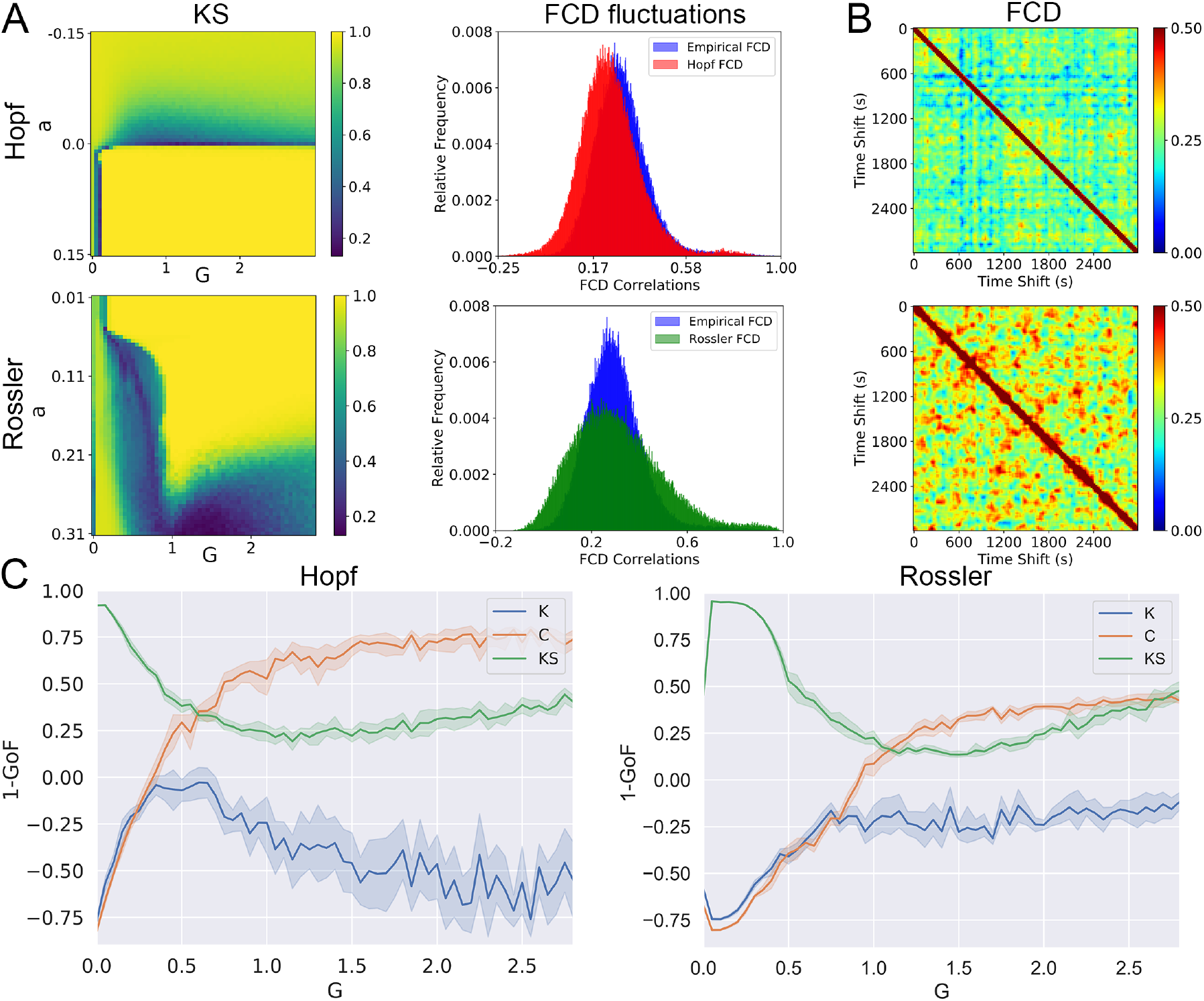
Whole-brain models fitted to dynamic observables. (a) KS distance between the distribution of simulated and empirical FCD values computed for all parameter combinations and for both models. Note that the best fit was obtained for *a* ≈ 0 for the Hopf model and for high *a* values for the Rossler model. The right panel displays the optimal distribution of FCD values for both models, together with the empirical distribution. (b) Examples of FCD matrices for both models computed using the optimal parameters obtained from the optimization of the KS distance. (c) KS distance, metastability (K) and synchrony (C) distance between simulated and empirical data as a function of *G*, computed for the best *a* obtained using the KS statistic.

Figure 4B shows for both models examples of simulated FCD matrices that minimized the KS distance to the empirical FCD. For the Rossler model, positive correlations were more widespread; in particular, the observation of positive correlations far from the main diagonal implied similar FC matrices for temporal windows at distant times. A similar pattern was observed in the example of the empirical FCD matrix provided in Fig. 2.

We computed the distance between simulated and empirical metastability and synchrony by subtracting the simulated values and normalizing, as explained in the Methods section. Fig. 4C shows how the KS distance, the metastability (K) and the synchrony (C) distance change as a function of *G* for the best parameter *a* obtained using the KS statistic. Neither model presented a minimum of the KS distance curve matching those of the other two metrics.

### D. Comparison of the optimal distance metric values between models

Figure 5 presents a comparison of the normalized distance (1 minus the goodness-of-fit, 1-GoF) for all the metrics and empirical observables reproduced by both models. These values were obtained from the exhaustive optimization procedures presented in Figs. 3 and 4. We assessed the effect size for the difference between the distributions of optimal distance metrics by means of Cohen’s d (Fig. 5, bottom panel)^70^, finding that all distance metrics had similar values for both models, with the exception of the Frobenius distance computed between simulated and empirical FC matrices, for which the Hopf model presented a significantly higher goodness of fit.

**FIG. 5.**
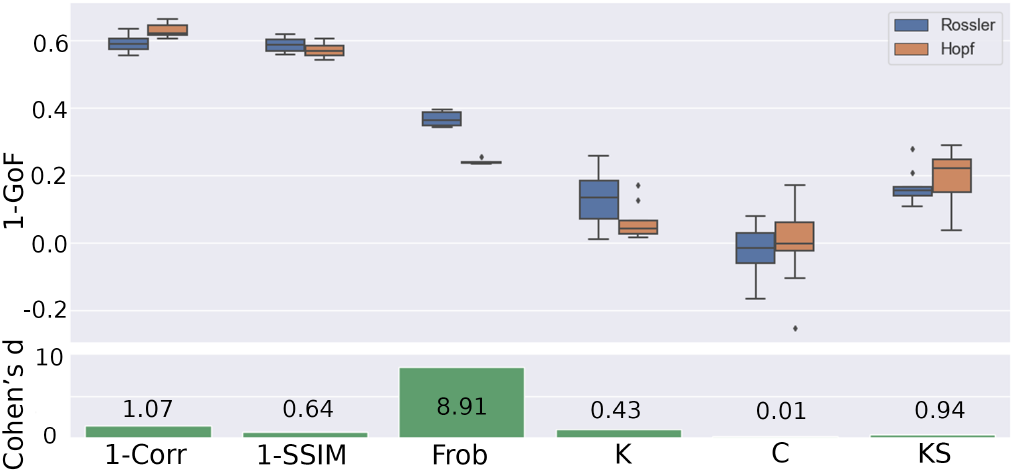
Comparison of 1 minus the goodness-of-fit (1 − *GoF*) for all metrics and observables between both models. The bottom panel presents the effect size for the difference in terms of Cohen’s d.

### E. Fitting multiple simultaneous observables

We investigated the capacity of each model to simultaneously reproduce multiple observables. For this purpose, we obtained the optimal parameters that reproduced a certain observable (following the exhaustive exploration procedure depicted in Figs. 3 and 4) and then compared how well the dynamics using those parameters could reproduce all other observables.

The results obtained from this analysis are presented in Fig. 6A. The value in a given matrix entry corresponds to the distance metric indicated in the column evaluated using the parameters that optimize the distance metric indicated in the row. For example, the value 0.28 in the second row and third column of the Hopf model matrix corresponds to the Frobenius distance between the empirical and simulated FC matrices obtained by simulating the Hopf model using the parameters that optimized the 1-SSIM. The bars on the right side of the matrices indicate the sum of values across columns, thus visualizing how well the parameters that optimize the distance metric in each row are capable of reproducing all other observables.

Figure 6B presents a graph representation of the generalization matrices in Fig. 6A. Each node represents a distance metric, and an arrow from one node to another indicates that the parameters optimizing the source metric result in low values for the target metric. For a given pair of metrics A and B, we computed the Cohen’s d between the optimal distance metric A values obtained after exhaustive exploration of the parameter space, and the distance metric B values obtained using the same parameters. A low Cohen’s d implies that both metrics yield similar values for the same model parameters; in other words, that those parameters simultaneously reproduce both observables. We computed the Cohen’s d for all pairs of metrics and then applied a threshold to display only the lowest 25 % Cohen’s d values as connections between the corresponding nodes.

**FIG. 6.**
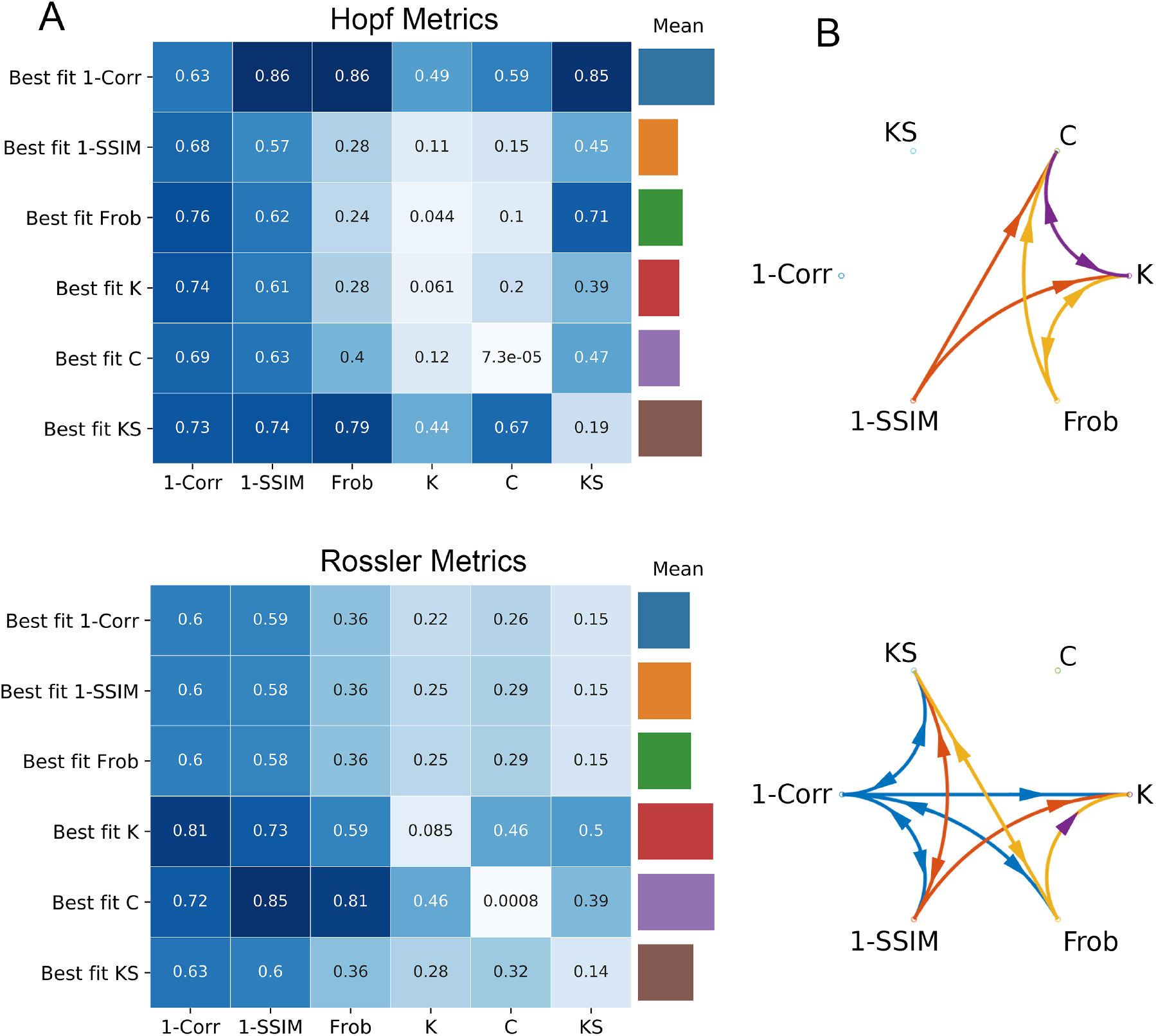
Generalization between empirical observables. (a) Matrix containing the distance metrics indicated in the columns evaluated using the parameters that optimize the distance metrics indicated in the rows. The bars on the right side of the matrix indicate the average across all columns. (b) Each distance metric is represented as a node in a graph where arrows indicate that the parameters optimizing the source metric result in low values for the target metric as well. For metrics A and B (represented by two nodes in the graph), we computed the Cohen’s d between the optimal distance metric A values obtained after exhaustive exploration of the parameter space, and the distance metric B values obtained using those same parameters. We applied a threshold to keep 25 % of the connections with the lowest Cohen’s d.

We can observe that the Hopf model optimized to reproduce *K* and *C* performed better than the Rossler model across all other metrics. On the other hand, for FC matrices compared using the 1-Correlation distance, and for FCD distributions compared using the KS distance, the Rossler model outperformed the Hopf model across all other metrics.

An important observation is that, in spite of its comparatively poor performance for *C* and *K*, the Rossler model optimized to reproduce the empirical dynamics encoded in the FCD matrix resulted in model parameters that also reproduced the “static” FC matrix, according to the three distance criteria (1-Correlation, 1-SSIM and Frobenius distance). Conversely, fitting the Rossler model to the “static” FC resulted in model parameters that approximated the empirical FCD distribution. This behavior was not seen for the Hopf model, for which the optimization of FCD resulted in parameters that failed to approximate the static FC, and vice-versa. In other words, the Rossler model did not force a choice between reproducing the FC over long temporal scales (FC matrix) and reproducing the FCD.

### F. Comparison of generalization performance between models fitted to FC and FCD

Figure 7 illustrates how the Rossler model is capable of simultaneously approximating static and dynamic FC, while the Hopf model produces inconsistent results when optimized to reproduce one of them and is then tested to reproduce the other. The first row of both left and right panels shows the optimal FC matrix (using the Frobenius distance) and the distribution of FCD values for the parameters that optimize the reproduction of the FC matrix. Conversely, the second row shows the FC matrix computed using the parameters that minimize the KS distance between empirical and FCD distributions, and then the distribution of FCD values using those parameters.

**FIG. 7.**
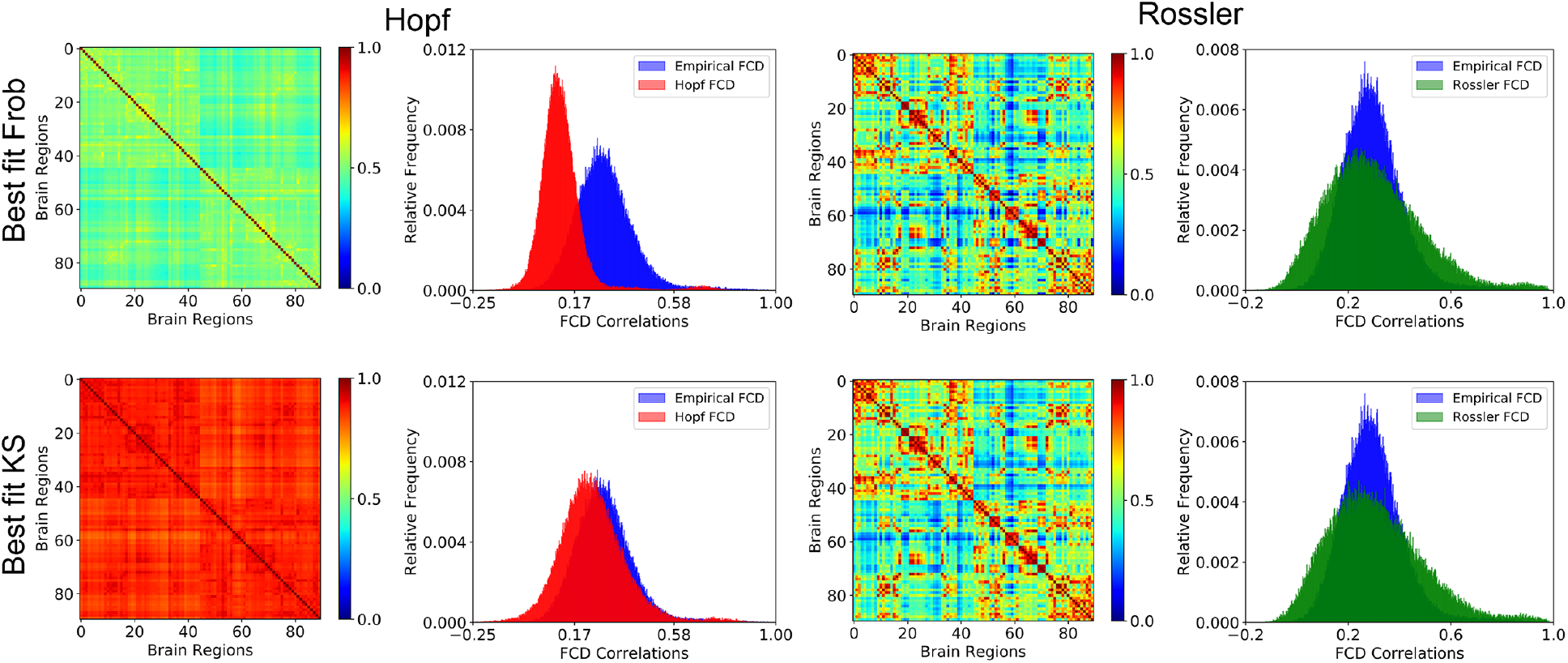
The Rossler model is capable of simultaneously approximating the empirical FC matrix and the distribution of FCD values. Each panel (left and right) contains in its first row the best reproduction of the FC matrix (using the Frobenius distance) and the distribution of FCD values for the parameters that optimize the reproduction of the FC matrix; conversely, the second row shows the FC matrix computed using the parameters that minimize the KS distance between empirical and FCD distributions, and then the distribution of FCD values using those parameters. The left panel illustrates the failure of the Hopf model to simultaneously reproduce both observables, which is achieved by the Rossler model, as shown in the right panel.

From the left panel, it is clear that the Hopf model tuned to reproduce the FC matrix grossly misrepresented the distribution of FCD values and vice-versa. As expected from the results presented in Fig. 6, the Rossler model simultaneously approximated the FC matrix and the distribution of FCD values with the same combination of parameters.

## IV. DISCUSSION

Speaking of Newton’s second law, the British astronomer Arthur Eddington once said: “*A force is whatever we need to put on left side of the equation to obtain results that agree with the observed motions*”^71^. In this view, physics is interpreted as an effort to produce increasingly more accurate models capable of describing the dynamics found in nature. Contemporary physics has become entangled with abstract mathematics to the point that formal structures are by themselves considered an important guide for theory building. In comparison, theoretical neuroscience is a much younger field still in need of exploring different mechanisms to explain the dynamics observed in experimental data^61^, a project having much in common with Eddington’s interpretation of Newton’s second law.

Ever since the discoveries of Hodgkin and Huxley, computational models of neural activity can be formulated with a high degree of biophysical realism^72^. Advances in microscopy and related technologies have contributed to unveil intricate networks of synaptic connections at the scale of single neurons, allowing to reconstruct the wiring of complete cortical columns^73^. Large collaborative efforts, such as the Human Brain Project, strive to combine this information to produce highly detailed simulations of small cortical regions^74^. As computational power is increased and the available maps of synaptic connectivity are expanded, it is expected that realistic models will begin to furnish predictions at the macroscopic scale, i.e. the scale investigated with neuroimaging tools such as fMRI, EEG and MEG. These predictions will have to be consistent with the phenomenological models that successfully described the dynamics and functional connectivity of large-scale brain activity. Thus, our investigation of the potential mechanisms underlying dynamics at this scale is motivated by the need to inform and constraint the development of more biophysically realistic models.

We explored two different mechanisms behind the complex spatiotemporal dynamics of resting state brain activity. As shown in Figs. 3 and 4, a model based on noise-driven multistability between equilibrium solutions (Hopf model) required fine-tuning of the bifurcation parameter to adequately reproduce different empirical observables. Conversely, a model based on deterministic chaos (Rossler model) reproduced these observables over a wider range of the model parameter, for which the local dynamics also exhibited positive Lyapunov exponents. This “stretching” of dynamical criticality finds a parallel in the substitution of a singular critical point by an extended critical-like region as a consequence of local dynamics being coupled by a complex hierarchical-modular network^75^.

This property is theoretically attractive, since models whose complex behavior depends on fine parameter tuning are unlikely to exhibit the robustness manifest in experimental brain activity recordings.

Computational models are developed and implemented with the purpose of capturing certain features of brain activity, which depend on the scientific question and its associated hypotheses. However, models can be difficult to interpret if they are inconsistent when attempting to reproduce multiple features at the same time. As shown in the left panel of Fig. 7, the optimal fit of the Hopf model to the FCD distribution results in parameters that produce a meaningless FC matrix, which is perhaps the most widespread summary statistic computed from resting state fMRI data^4^. Ideally, phenomenological models of whole-brain activity should be capable of approximating both the “static” and dynamic FC; however, this could be difficult for noise-driven equilibrium models which require fine-tuning to dynamical criticality to produce rich temporal FC fluctuations. In the Rossler model we found a “stretching” of the range of model parameters that simultaneously fitted multiple empirical observables. This result suggests that the dichotomy between reproducing static or dynamic FC features could be avoided by the introduction of deterministic chaos in the local dynamics.

Models based on noise-driven multistability have found widespread applications in computational neuroscience^76^. Since their dynamics can be understood in terms of attractors connected by noise-induced transitions, these models are easier to interpret and construct with the purpose of producing certain predefined behaviors. Within the specific context of whole-brain models, the interplay between noise and deterministic equilibrium dynamics is sufficient to generate ongoing dynamics capable of exploring the repertoire of potential brain configurations^7^. The noise-driven exploration of this repertoire produces the dynamic fluctuations in functional connectivity that have been robustly established using several imaging modalities^23,49,77^. In general, the qualitative global behavior of whole-brain models can be understood in terms of the bifurcation diagrams of the local dynamics, at least in the case of weak coupling. Conversely, while coupled chaotic systems can exhibit multistable swiching by mechanisms such as chaotic itineracy^53^ and heteroclinic cycling^52^, local dynamics cannot be easily understood by bifurcation analysis due to the presence of strange attractors.

The inclusion of noise should not be disregarded as an *ad hoc* mechanism required to produce interesting dynamics. Instead, it should be considered as the potential manifestation of biological and physical processes occurring at multiple spatial and temporal scales. Neural systems are subject to a number of noise sources: the activity of a population of neurons embodied in the brain inevitably occurs in the presence of stochastic fluctuations due to thermal energy, ion channel chattering, intermittent neurotransmitter release, and irregular synaptic inputs from other neurons, among other sources of noise^76,78^. However, the biological interpretation of additive noise terms in large-scale models of whole-brain activity remains unclear. Following the principle of economy of explanation^79^, unknown sources of variability should be considered less satisfactory than endogenous variability produced by intrinsically non-equilibrium dynamics. Even though deterministic chaos may be appealing for large-scale phenomenological models, it is less clear how to systematically construct biophysically realistic models whose variability stems from chaotic dynamics, and whose output is consistent with experimental data.

Another relevant point is whether empirical data supports the presence of chaos in brain dynamics. This is a contentious issue, possibly because current experimental tools are insufficient to produce evidence that can be considered as definitive. Over the past decades, several studies reported the presence of chaotic dynamics in time series obtained from an ample variety of neural systems^59,60^; however, an equally large literature has been published arguing against this possibility^80,81^. Theoretically, it is accepted that the inherent instability of chaotic dynamics facilitates the extraordinary ability of neural systems to respond quickly to changes in their external inputs^82^, to flexibly transition between behavioral patterns as a consequence of environmental changes, and to explore the large repertoire of dynamical states that endows neural circuits with remarkable computational capabilities^83^. Chaotic dynamics in the brain could emerge in several ways, such as from intrinsic mechanisms within individual neurons^84^, or from the collective dynamics of neural networks^85–87^. While our results do not demonstrate the presence of deterministic chaos in large-scale brain activity, they illustrate how even the simplest model of coupled chaotic oscillators is capable simultaneously reproducing “static” and dynamic FC observables. This result should be taken into consideration by future model building efforts, independently of the deeper question of whether chaotic dynamics represents an intrinsic feature of brain activity.

It must be stressed that noise and deterministic chaos are not mutually exclusive mechanisms to produce multistable brain dynamic. As recently shown by Orio et al.^88^, moderate noise can enhance the multistable behavior of chaotic neural networks, resulting in an ampler exploration of the synchronization repertoire, with very high levels of noise eventually abolishing multistability. While this study investigated a conductance-based neural model wired with a small-world network topology, presumably noise-enhanced multistable chaotic dynamics can also be found in Rossler oscillators coupled by realistic anatomical connectivity, a possibility that should be addressed by future studies.

Our study presents some limitations pointing the way towards future improvements. First, while we considered instantaneous interactions between the network nodes, the interplay between noise and conduction delay is a key factor to reproduce the dynamics of spontaneous brain activity fluctuations^89^. However, the combination of slow temporal sampling by fMRI and fast conduction speeds through long-range myelinated axons likely attenuates the effects of omitting delays in our models. Second, we did not obtain the anatomical connectivity networks from the same subjects who were recorded with fMRI. Instead, DTI recordings were obtained from another group of healthy participants who can be considered as representative of an adult population. However, individual connectivity matrices should be used when modeling the large-scale dynamics of individuals who could present structural abnormalities as a consequence of neurological or psychiatric impairments. Finally, we focused on modeling unusually long recordings (50 minutes) of awake subjects, which contributed towards more robust estimates of dynamic FC measures. However, the biological implications of our findings should be explored by investigating subjects during other states of consciousness, e.g. states of such as deep sleep, general anaesthesia, or in patients diagnosed with disorders of consciousness^90^. It is tempting to speculate that the level of consciousness will be paralleled by the degree of chaoticity in the best fitting model, a possibility that will be explored in future studies.

In conclusion, we showed that chaotic dynamics give rise to some interesting features in whole-brain activity models, out-performing noise-driven equilibrium models in the simultaneous reproduction of multiple empirical observables. While simplified phenomenological models may appear to be overly detached from the intricate details of the human brain, they are nevertheless important to propose conceptually simple mechanisms that more realistic models should eventually strive to reproduce. Our results identified some attractive features of deterministic chaos that should not be neglected by future modeling efforts, even in the simultaneous presence of noisy inputs. Facing up to the challenge of imprinting and interpreting deterministic chaos in more realistic whole-brain models could be a key to understand how the human brain is capable of producing an ever-changing stream of complex activity patterns.

## ACKNOWLEDGMENTS

This work was supported by funding from Agencia Nacional De Promocion Cientifica Y Tecnologica (Argentina), grant PICT-2018-03103. The authors acknowledge the Toyoko 2020 program for granting cloud computing services.

